# Automated Reconstruction of a Serial-Section EM *Drosophila* Brain with Flood-Filling Networks and Local Realignment

**DOI:** 10.1101/605634

**Authors:** Peter H. Li, Larry F. Lindsey, Michał Januszewski, Zhihao Zheng, Alexander Shakeel Bates, István Taisz, Mike Tyka, Matthew Nichols, Feng Li, Eric Perlman, Jeremy Maitin-Shepard, Tim Blakely, Laramie Leavitt, Gregory S.X.E. Jefferis, Davi Bock, Viren Jain

## Abstract

Reconstruction of neural circuitry at single-synapse resolution is a key target for improving understanding of the nervous system in health and disease. Serial section transmission electron microscopy (ssTEM) is among the most prolific imaging methods employed in pursuit of such reconstructions. We demonstrate how Flood-Filling Networks (FFNs) can be used to computationally segment a forty-teravoxel whole-brain *Drosophila* ssTEM volume. To compensate for data irregularities and imperfect global alignment, FFNs were combined with procedures that locally re-align serial sections as well as dynamically adjust and synthesize image content. The proposed approach produced a largely merger-free segmentation of the entire ssTEM *Drosophila* brain, which we make freely available. As compared to manual tracing using an efficient skeletonization strategy, the segmentation enabled circuit reconstruction and analysis workflows that were an order of magnitude faster.

## Introduction

The description of neural function in terms of the circuit architecture of individual neurons and their connections remains a fundamental goal in neurobiology (Shepherd and Grillner 2018). Progress towards this goal has recently been enabled by advances in “connectomics”, specifically nanometer-resolution volume imaging as well as computational methods for visualizing and annotating 3d image data. Scale, however, remains a fundamental constraint: most connectomic studies have been limited to imaging volumes that are millions of cubic microns or less, and in many cases only a fraction of the imaged data have been reconstructed and analyzed (Kornfeld and Denk 2018). In practice this has meant limiting connectomics studies to, for example, fractions of mouse (Bock et al. 2011; Helmstaedter et al. 2013; Kim et al. 2014; Kasthuri et al. 2015; W.-C. A. Lee et al. 2016), rat (Schmidt et al. 2017), rabbit (Anderson et al. 2011), songbird (Kornfeld et al. 2017), zebrafish (Wanner et al. 2016), or *Drosophila (Takemura et al. 2013; Takemura, Aso, et al. 2017; Takemura, Nern, et al. 2017; Shan Xu et al. 2020)* brains, or else characterizing complete but smaller brains such as *C. elegans* (White et al. 1986; Varshney et al. 2011) or larval *Drosophila* (Ohyama et al. 2015).

A recent imaging milestone provides both completeness and significantly increased scale compared to previous synapse-resolution connectomic studies: a serial section transmission electron microscopy (ssTEM) volume of a complete adult *Drosophila* brain imaged at 4×4 nm resolution and sectioned at 40 nm thickness, known as the “full adult fly brain” (FAFB) dataset (Zheng et al. 2018). This trove of image data has the potential to reveal fundamental aspects of the structure and function of the *Drosophila* central nervous system, provided that significant hurdles related to the manual and automated reconstruction of the neurons and their connections can be mitigated or overcome (Funke et al. 2016).

For example, a complication of ssTEM and serial section microscopy in general is the necessity and difficulty of computationally aligning independently imaged sections into a coherent three-dimensional volume, in which physical structures in the tissue are represented at stable XY locations across consecutive imaging planes. In fact, large serial section electron microscopy datasets often require warping, or elastic alignment algorithms (Saalfeld et al. 2012), as well as specific compensation for data irregularities that invariably occur such as tears, folds, cracks, and contaminant particles (Wetzel et al. 2016; Zheng et al. 2018). For large datasets, complete identification and compensation for artifacts is historically intractable, with some misalignments generally persisting throughout subsequent analysis efforts. When tracing over these misalignments, there is a significantly increased danger of introducing errors such as mergers (in which two or more processes are erroneously connected to one another), for both automated algorithms as well as trained human annotators (Schneider-Mizell et al. 2016).

In order to compensate for imperfections in global alignment of the full adult fly brain (FAFB) ssTEM data, we integrated a new “local realignment” procedure into the flood-filling network (FFN) segmentation method (Michał Januszewski et al. 2018), which operates on local subvolumes as an intermediate step towards building a whole volume segmentation. Local realignment improved the alignment quality of subvolumes provided to the FFN, and when combined with a procedure that further gated FFN segmentation depending on resulting data quality, reduced total merge errors by an order of magnitude (Fig. 3). In an additional “irregular section substitution” process, we automatically detected data irregularities that persisted after local realignment and, where possible, replaced affected regions with data from neighboring sections. This reduced split errors (where two processes are erroneously disconnected from one another) and thus tripled the “expected run length” (Michał Januszewski et al. 2018) (Fig. 4). Finally, for severe situations in which multiple sections of raw data were missing, we used a Segmentation-Enhanced CycleGAN (SECGAN) to synthesize replacement imagery (Michal Januszewski and Jain 2019).

**Figure 1.**
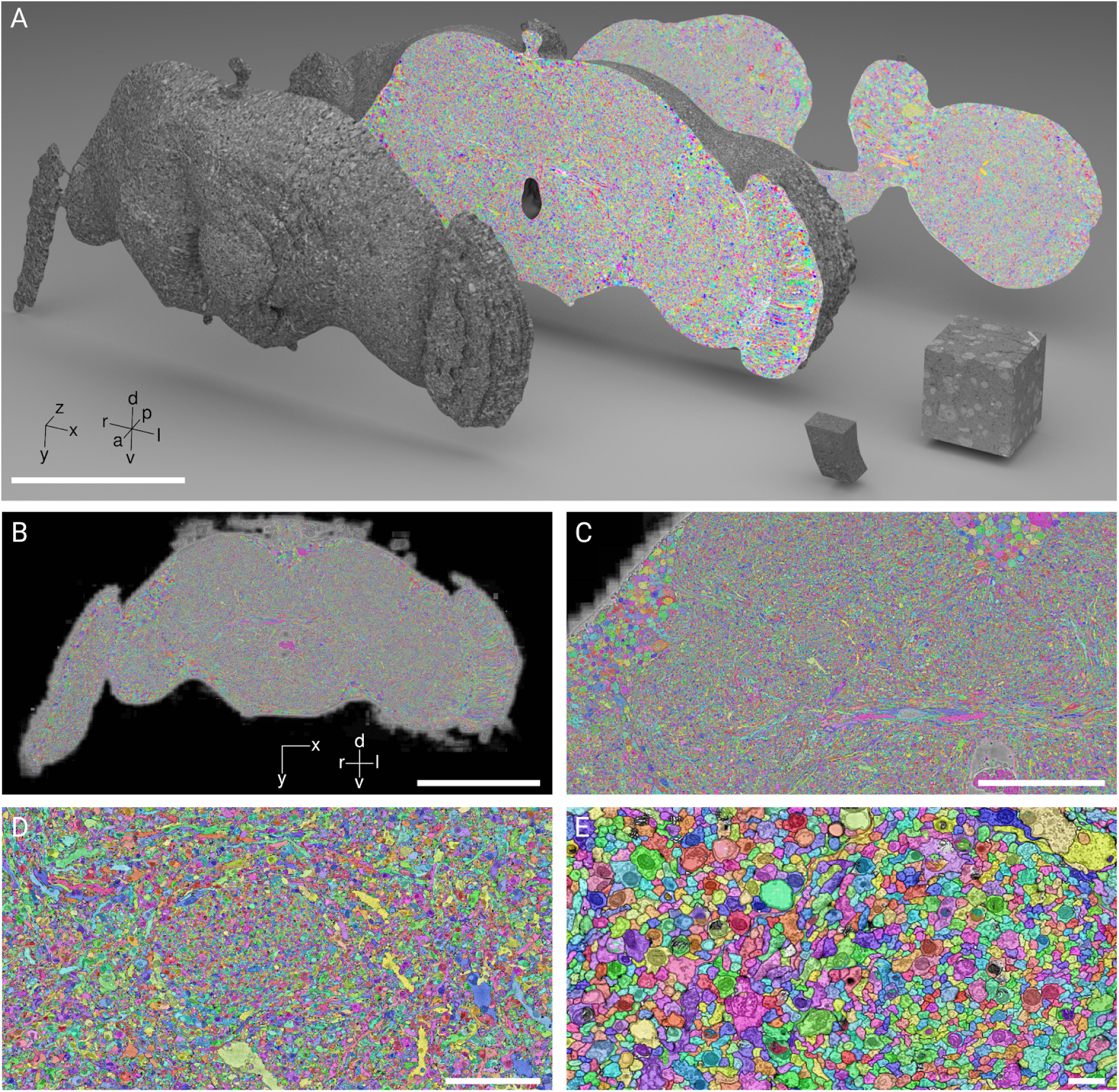
Dense segmentation of the entire fly brain via FFNs. **(A)** 3D rendering of a smoothed tissue mask of the FAFB dataset. Arbitrary coronal sections (dataset XY plane) reveal the FAFB-FFN1 segmentation throughout the interior. In the lower right, two recent dense segmentation benchmarks, from the songbird tectum (Michał Januszewski et al. 2018) and *Drosophila* optic lobe (Takemura et al. 2015), for comparison. Scale bar 200 μm. Exterior surface of FAFB is pseudo-textured. **(B-E)** Increasing zoom levels of coronal sections intersecting the mushroom body peduncular tract (right hemisphere), FAFB-FFN1 segmentation overlaid on CLAHE processed raw data. Scale bars 200, 50, 10, 1 μm. d, dorsal; v, ventral; a, anterior; p, posterior; r, right hemisphere; l, left hemisphere.

**Figure 2.**
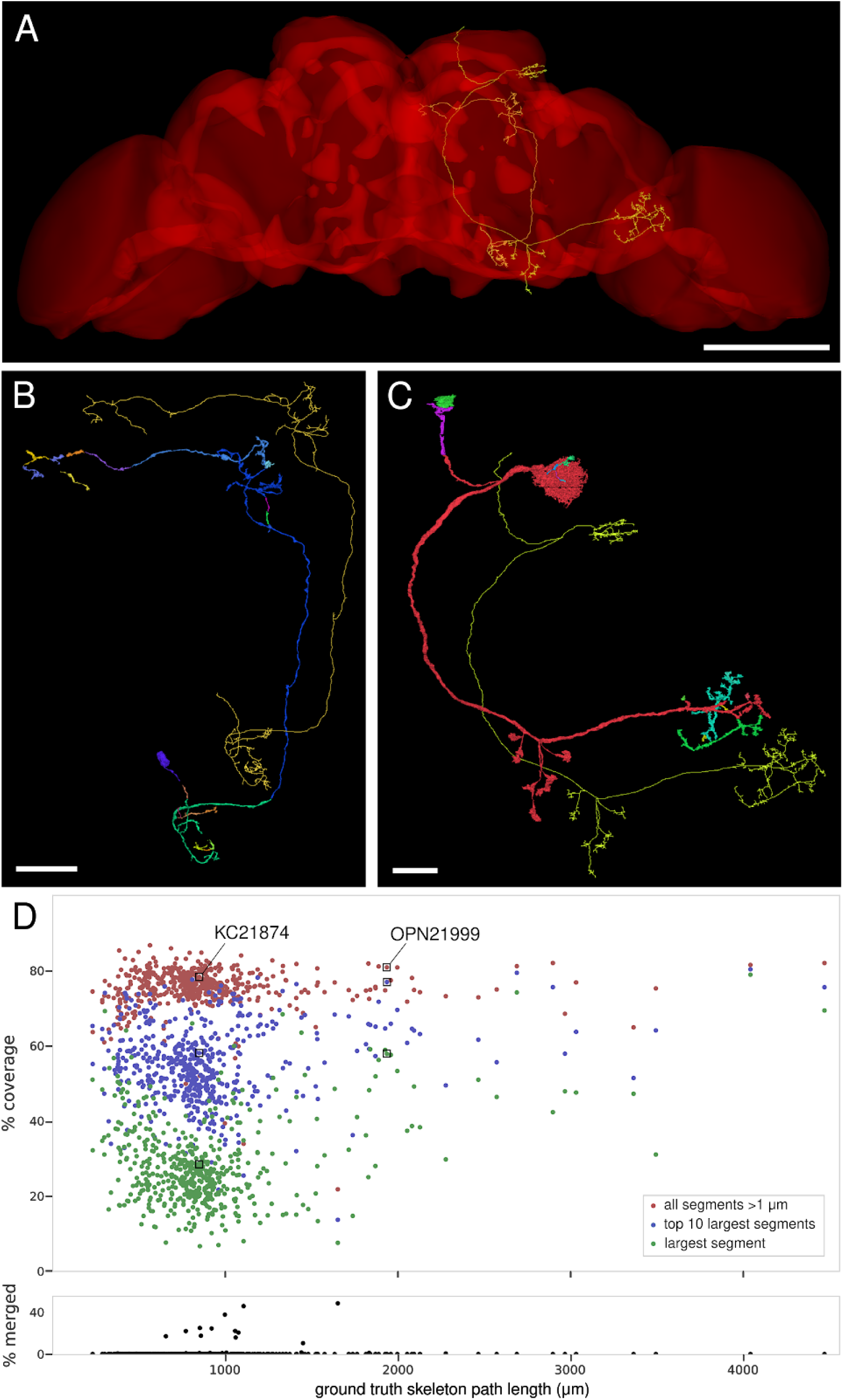
Automated neuron reconstructions validated against manual neuron tracings. **(A)** Manual skeleton tracings of Kenyon cell (KC) 12874 and olfactory projection neuron (OPN) 21999 (Zheng et al. 2018), overlaid on the reference fly brain (Jefferis 2018a). Scale bar 120 μm. **(B)** The same KC12874 skeleton as in (A), displayed alongside the 23 FFN segments that overlap with it for more than 3 μm of path length (offset for clarity, segments colored arbitrarily). FFN segments cover significant skeleton path length without erroneously merging into any neighboring neurites. Scale bar 20 μm. **(C)** As (B), for the 14 segments that overlap OPN21999. **(D)** Top, for each ground truth skeleton (n=525), three points showing coverage of the skeleton by FFN segments as a percent of total path length. The point groups are: just the largest segment (by path length covered) for each skeleton (green), the summed path length covered by the five largest segments per skeleton (blue), or the summed path length per skeleton for all segments greater than 1 μm (red). Points for KC12874 and OPN21999 are indicated. Bottom, for each skeleton, the percent of total path length covered by segments that erroneously merge multiple skeletons. For 478 out of 525 cells there are zero merge errors, and for 509 cells the merged path length is < 1%.

**Figure 3.**
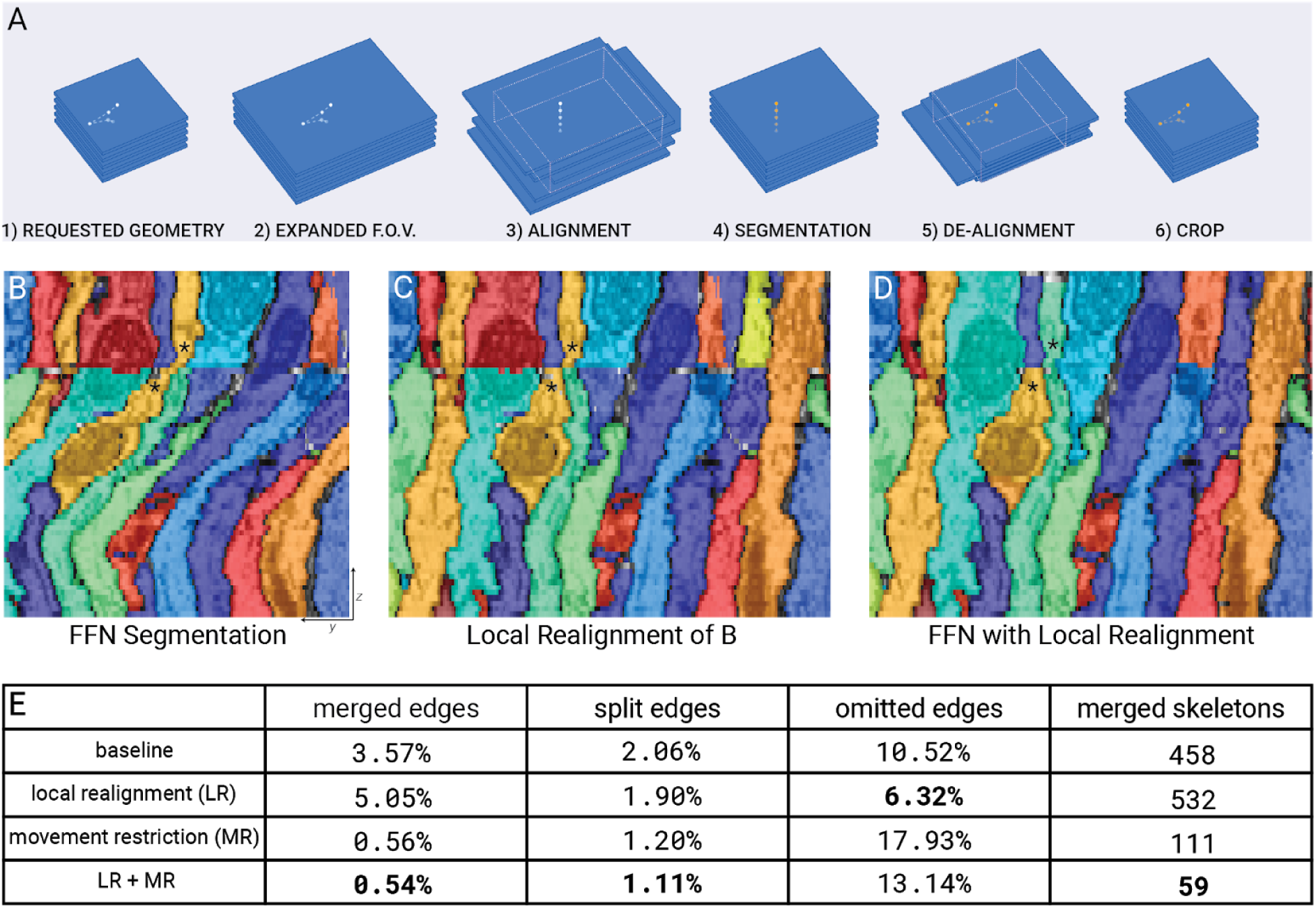
Local realignment (LR). **(A)** LR procedure; a local subvolume is requested for FFN inference (1-2); this is aligned according to section image content (3) and then cropped and segmented (4); the segmentation output is then dealigned (5) and cropped back to the original requested subvolume dimensions for incorporation into a coherent output volume (6). The expansion of the subvolume bounds in (2) is computed to account for the two subsequent rounds of cropping. **(B)** YZ view of FFN segmentation at for example location in FAFB without LR. **(C)** The same segmentation result as (B), with LR applied *post-hoc*. This reveals more clearly the impact of the discontinuity in section alignment, including many clear split errors as well as a merge error (stars). **(D)** Segmentation of the same location as (B) and (C), now with LR applied as a preprocessing step prior to segmentation. This fixes the merge error as well as most splits. **(E)** Impact of LR on skeleton metrics for 16×16×40 nm resolution FFN segmentation of the Sample E mushroom body cutout. LR fixes many merge and split errors, but cannot correct heavily distorted areas or other types of data irregularity such as damaged or occluded sections. In combination with movement restriction to address these other cases, LR reduced merge errors by an order of magnitude while minimizing splits and omitted areas.

**Figure 4.**
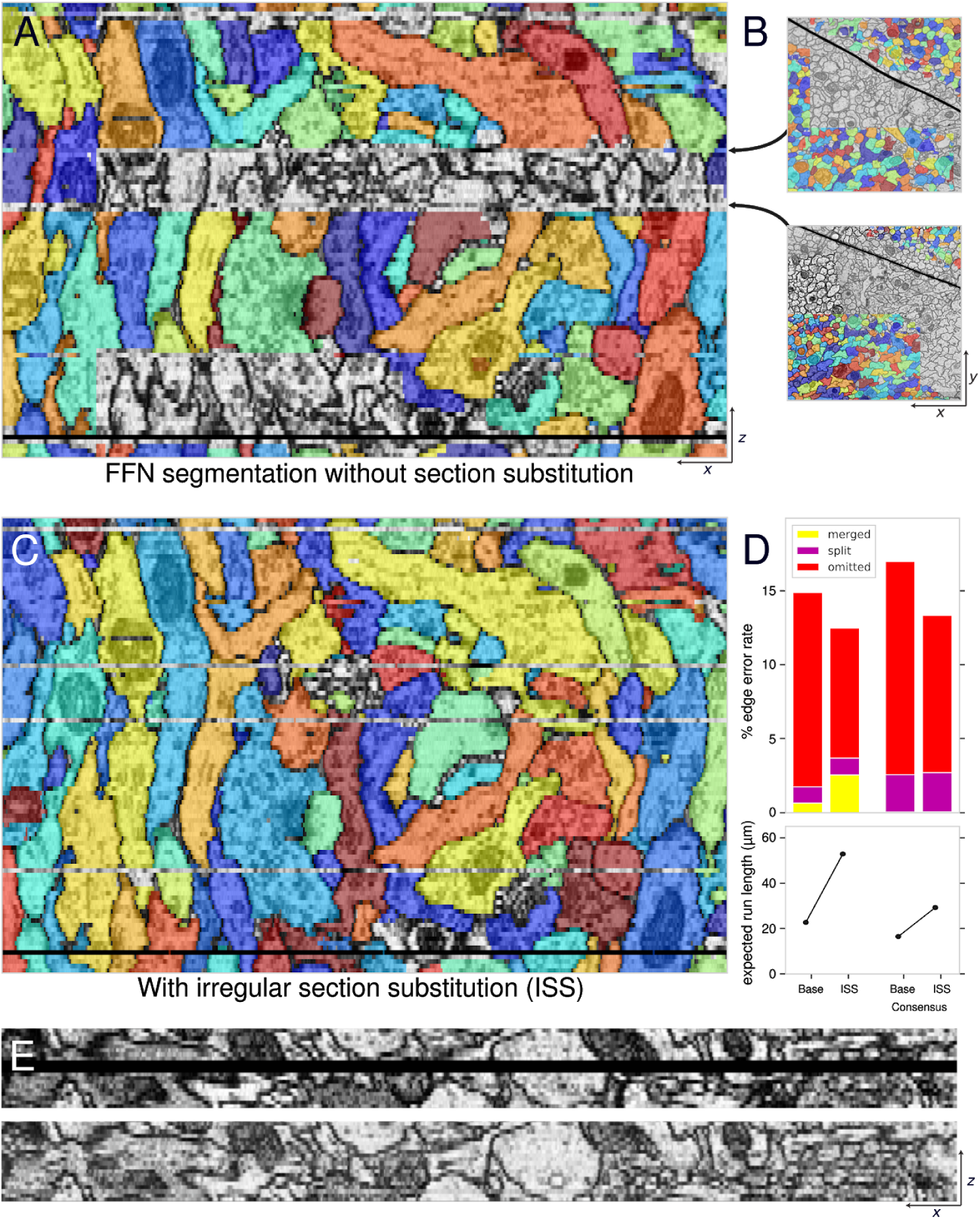
Irregular section substitution (ISS) and SECGAN. **(A)** XZ view of FFN segmentation for an example location in FAFB, with LR and movement restriction preprocessing (Fig. 3). Segments are interrupted at multiple sections due to data artifacts incompletely addressed by LR. When artifacts occur on nearby sections, movement restriction can cause the whole area to be left unsegmented. **(B)** XY views of two sections from (A) showing tissue fold artifacts. **(C)** The same view as (A), but with automatic ISS applied prior to segmentation. **(D)** Impact of ISS on skeleton metrics for 16×16×40 nm resolution FFN segmentation of Sample E. Top, percent remaining skeleton edge errors by subtype. Bottom, expected run length, the average error-free path length of segments under uniform sampling along skeletons. ISS substantially reduces omission errors and nearly triples expected run length, but also causes some new merge errors. However, the increased merge error effect is nullified by later oversegmentation consensus procedure (Fig. 5). **(E)** Top, XZ view of three consecutive missing sections (Z indices 3595-3597). Bottom, SECGAN interpolation replaces the missing sections with synthesized imagery.

In the following we (1) describe a multi-scale FFN segmentation pipeline for FAFB, (2) show how local realignment and irregular section substitution can be used to deal with imperfect alignment and artifacts, (3) characterize reconstruction accuracy using ground truth skeletons, and (4) show how the automated segmentation results can be used to assist and accelerate manual reconstruction and analysis of neural circuits. We provide the complete segmentation and accompanying metadata such as derived skeletons as a public resource to assist further efforts related to FAFB, *Drosophila* connectomics, and connectomic algorithm development.

## Results

### Flood-filling segmentation of FAFB

The primary result presented here is an automated segmentation of neuronal processes densely covering the entire FAFB dataset (Fig. 1), which contains 40 teravoxels of tissue within a 995×537×283 μm EM volume resulting from a correlation- and feature-based deformable alignment of ∼21 million raw ssTEM camera images (Zheng et al. 2018; Khairy, Denisov, and Saalfeld 2018). We segmented FAFB via a multistage pipeline whose workhorse was the FFN (detailed below), operating in a coarse-to-fine mode that reduced computational cost by an order of magnitude.

The segmentation result, referred to as “FAFB-FFN1”, is largely an oversegmentation, in which each neuron in the dataset corresponds to multiple automated segments. Conversely, merge errors, in which a single segment overlaps multiple neurons, are rare. We evaluated these properties by comparing the segmentation to a set of 525 ground truth neuronal skeletons produced by human tracers (Zheng et al. 2018, 2020) (Fig. 2). For each ground truth skeleton, we found all FFN segments that overlapped skeleton node positions; ground truth skeletons could then be visualized alongside their overlapping segments to assess the quality of segmentation coverage for each neuron (Fig. 2B-C). We also checked whether each segment overlapped only a single skeleton, versus erroneously merging multiple skeletons together. Finally, for each ground truth skeleton we analyzed the percent of total path length correctly covered by merge-free segments versus the percent impacted by merge errors (Fig. 2D). Out of 525 ground truth skeletons, 509 had negligible merge errors.

We further quantified FAFB-FFN1 final segmentation quality using summary metrics based on the ground truth skeletons (Michał Januszewski et al. 2018). *Edge accuracy* categorizes skeleton edges as either correctly reconstructed, or else falling into one of three error categories: split, omitted, or merged. FAFB-FFN1 achieves edge accuracies of 91.5% correct, 4.4% split, and 3.3% omitted, with only 0.82% merged. *Expected run length (ERL)*, computes the expected error-free path length in the segmentation given a uniformly sampled starting point along a ground truth skeleton. FAFB-FFN1 achieves an ERL of 199 μm, comparable to the path error rates computed between multiple expert human tracers in adult (61 μm / error) and larval (27 μm / error) *Drosophila* brain volumes (Zheng et al. 2018; Schneider-Mizell et al. 2016). We also used the skeleton metrics extensively to evaluate the quality of intermediate results throughout the development of the detailed FAFB-FFN1 pipeline described below (Figs. 3-5).

### Handling input data irregularities

To segment FAFB, it was critical to address misalignments and data irregularities such as damaged, occluded, missing, or distorted sections. While these issues affect most connectomic datasets, they are particularly prevalent in serial-section data. FFN segmentation without specific procedures to address irregularities resulted in unacceptably high error rates, particularly for merge errors (Fig. 3). One strategy to handle irregularities is simply to prevent the FFN from moving into any field of view where irregularities are detected (Michał Januszewski et al. 2018). This reduces merge errors while increasing splits. More critically, it can also result in significant omission errors, where regions of the dataset are left unsegmented (Figs. 3A, 4A). Therefore, we addressed these issues more directly via three “preprocessing” steps (attempting to correct issues in the raw inputs before they are passed to the FFN): *Local Realignment (LR), Irregular Section Substitution (ISS), and Segmentation-Enhanced CycleGAN (SECGAN)*.

LR sidesteps the difficulties inherent in globally aligning large serial-section datasets by correcting residual misalignment within each local subvolume block just prior to segmentation (Fig. 3A). We first automatically estimated residual misalignment in the globally aligned dataset via neighboring section cross-correlation template matching (see Methods). The resulting section-to-section shift estimates allowed each local subvolume block to be realigned on-the-fly prior to FFN segmentation, with additional raw image context drawn as needed per section from the underlying dataset. LR significantly improved segmentation, correcting many merge errors and splits (Fig. 3B-E).

ISS was also applied on-the-fly prior to FFN segmentation, to address the problems of damaged, occluded, missing, and distorted areas by selectively replacing these areas with data from neighboring sections (Fig. 4). We again used section-to-section cross-correlation, now to detect all classes of irregularity remaining after LR (e.g. Fig. 4A-B). We then evaluated, for each detected irregular section, whether the cross-correlation would be improved within the local subvolume block by replacing the irregular section with a copy of the previous neighboring section. Accepting substitutions that passed this evaluation and applying them prior to FFN segmentation significantly improved contiguity and completeness of the result (Fig. 4C), nearly tripling expected run length (Fig. 4D). While ISS did incur an increase in merge errors, this negative impact was effectively nullified by the later oversegmentation consensus stage (see below). For any areas where detected irregularities persisted after both LR and evaluation for ISS, we fell back to simply preventing the FFN from moving into the area. This corrected some remaining merge errors, at the cost of omission and split errors that were addressed via later fill-in and agglomeration stages (see below).

Finally, in three locations in the volume where multiple consecutive sections were partially or completely missing, we trained a SECGAN to synthesize (i.e., “hallucinate”) missing data (Fig. 4E). The SECGAN enabled the FFN to trace through about 60% of the neurites in such locations, on average.

### Overall FAFB segmentation pipeline

FFN segmentation was first performed at reduced resolution (16×16×40 nm) to efficiently handle large structures at the scale of the whole fly brain, before switching to higher resolutions to fill in remaining gaps and capture finer structures (Fig. 5A). As a result, the FFN considered 2.2 teravoxels of tissue for each complete run at 16×16×40 nm resolution, but only 1.5 teravoxels of remaining unlabeled tissue at 8×8×40 nm (versus 8.8 teravoxels total) and only 4.2 teravoxels of remaining unlabeled tissue at 4×4×40 nm (versus 35.2 teravoxels total). Furthermore, resolution was selectively reduced within, but not across, tissue sections (i.e. in the X and Y, but not Z dimensions). This rendered most pipeline inputs more isotropic than the raw data, allowing network architectures previously developed on more isotropic datasets to be successfully redeployed.

**Figure 5.**
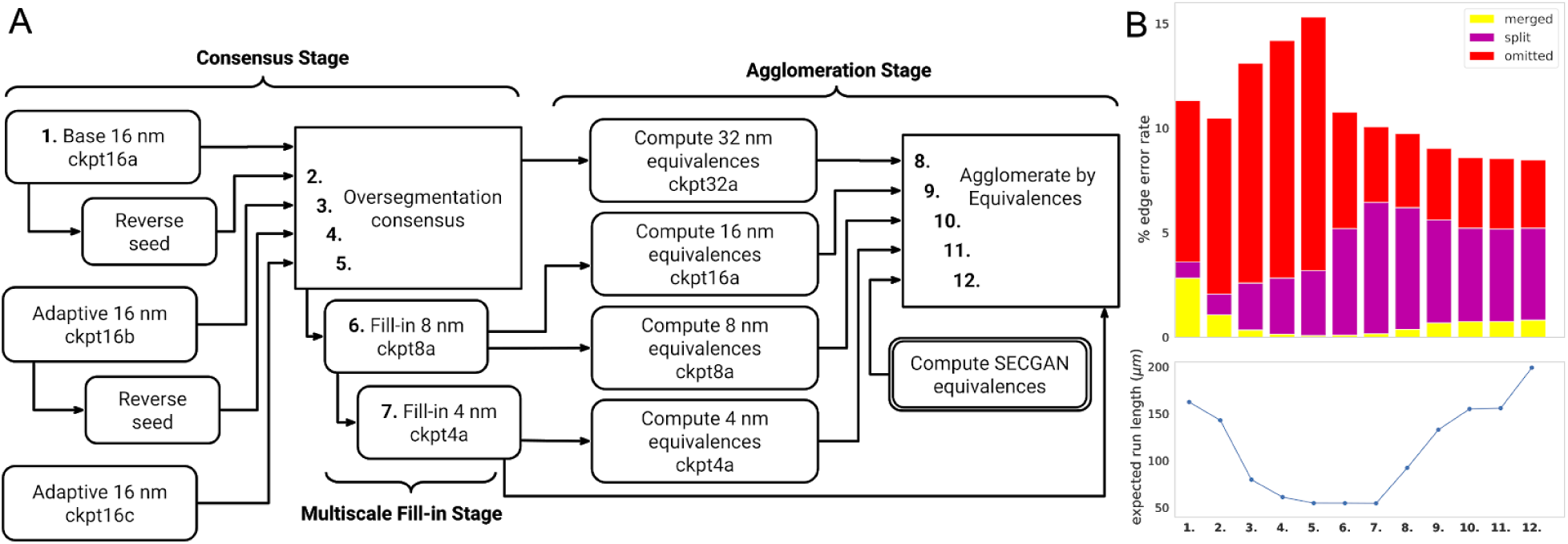
Overall FAFB-FFN1 segmentation pipeline. **(A)** Detailed pipeline steps and major stages, bold numerals highlight selected intermediate volumes in the pipeline used for evaluations in (B). Rounded boxes indicate FFN inference stages. Each FFN step indicates the model checkpoint used. **(B)** Evaluation metrics relative to ground truth skeletons at progressive stages of the pipeline, columns corresponding to pipeline stages with bold numeral labels in (A). Top, percent remaining skeleton edge errors by subtype. Bottom, expected run length.

The full segmentation pipeline consisted of three major stages: *consensus, fill-in*, and *agglomeration* (Fig. 5A). Each stage used the FFN in a different mode, and primarily addressed a different class of segmentation error (merges, omissions, and splits) as revealed in the skeleton metric evaluations (Fig. 5B). Every step involving FFN evaluation (Fig. 5A, rounded boxes) also included LR and ISS preprocessing, except the SECGAN computations which handled misalignment and data irregularity separately.

In the initial stage, we ran the FFN multiple times at 16×16×40 nm resolution from different starting conditions to produce five separate segmentations of the entire FAFB volume. We then combined these results via an “oversegmentation consensus” procedure that breaks segments at any location where the input segmentations disagree (Michał Januszewski et al. 2018). The resulting consensus segmentation has a very low merge error rate at the cost of accumulating split errors (Fig. 5B), as merge errors are retained only when present in *all* input segmentations, whereas split errors in *any* input carry through.

In the second stage, the FFN was only allowed to fill-in remaining empty areas while leaving pre-existing consensus segments unchanged, thus reducing omission errors without introducing significant new merge errors. The primary cause of omissions was inadequate image resolution; FFNs trained and run at 16×16×40 nm resolution were unable to follow some finer structures, leaving these regions empty. FFNs trained and run at 8×8×40 or 4×4×40 nm were able to fill in most of these areas (Fig. 5, steps 6-7).

In the third stage, we used the FFN to evaluate whether selected abutting segments from prior fill-in results should be merged together (i.e. automatically agglomerated). For each evaluation, we extracted small subvolumes (60×60×30 voxels) surrounding a local interface between two candidate segments from both the image volume and the segmentation, seeded the FFN from several points on each side of the interface, and accepted a merge for the segments if the FFN was able to flood significantly and consistently across the interface from both directions (Michał Januszewski et al. 2018).

We evaluated agglomeration candidates at multiple resolutions. For segments in the 16 nm segmentation (Fig. 5A, step 5) we evaluated candidates using an FFN trained and run at 32×32×40 nm. Segments in the 8 nm segmentation (Fig. 5A, step 6), including those created in consensus and fill-in stages, were agglomerated using FFNs trained at both 16×16×40 and 8×8×40 nm resolution. Segments in the 4 nm segmentation (Fig. 5A, step 7) were agglomerated at 4 nm. The automated agglomeration process corrected 30% of the remaining splits in the segmentation (1.5% of total edges), and was tuned to avoid committing significant new merge errors (Figure 5B).

Finally, we used SECGAN synthesized imagery in three regions of the volume to bridge multiple consecutive missing or irregular sections. Without SECGAN replacement, these areas split crossing processes for most of the neurons across the entire XY extent of the volume. We analyzed FFN segmentation of the SECGAN replacement imagery to agglomerate non-SECGAN segments across the gaps, repairing 150,000 splits and improving final ERL from 156 to 199 μm (Fig. 5B).

### Application of automated segmentation to circuit tracing workflows

Experiments with human tracers demonstrated the benefit of using FFN segmentation to assist typical biological analysis workflows. We assessed raw speed-up of neuron tracing in several brain areas (Fig. 6) as well as benefits for trans-synaptic circuit tracing efforts (Fig. 7). Some neuron tracing experiments (Fig. 6) were conducted using earlier preliminary versions of the FFN segmentation, with ERLs of 40-50 μm (see Methods); because of the human effort involved these were not repeated using the more complete final FAFB-FFN1. In some cases, tracers worked directly with FFN segments, while for other workflows we first automatically skeletonized the segmentation via blockwise TEASAR (Sato et al. 2000) and the tracers then worked with the skeleton fragments. Depending on the specific workflow, observed benefits ranged from several-fold to more than an order of magnitude in human effort saved.

**Figure 6.**
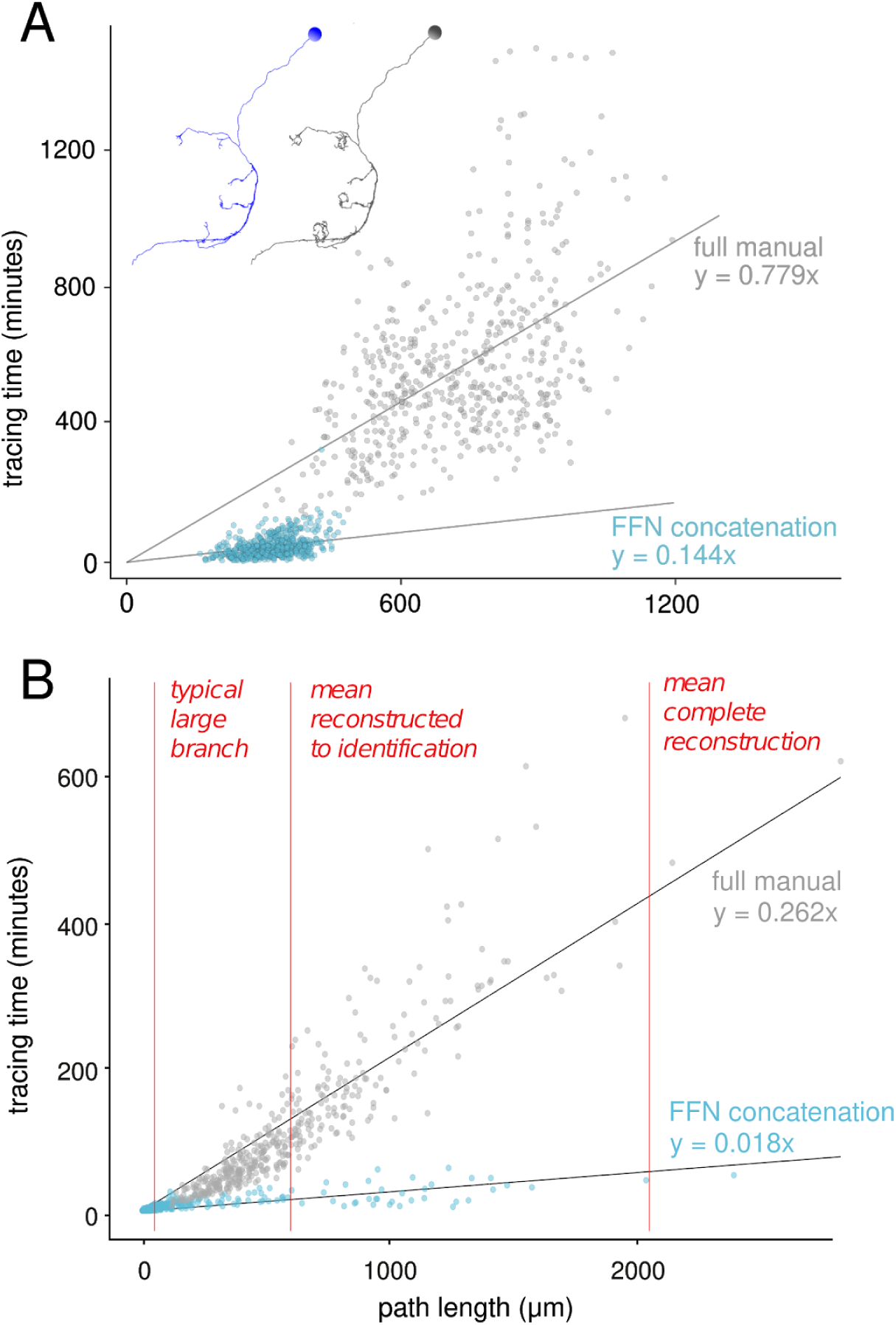
Segmentation-assisted neuron tracing. **(A)** Tracing speed (person-minutes per μm path length) for Kenyon cells in the mushroom body calyx. Cells are grouped according to the reconstruction methodology: “full manual” (gray points, n=545) or “FFN concatenation” (blue points, n=601), in which FFN segmentation-derived skeleton fragments were linked together. Inset illustrates for one cell the targeted level of reconstruction completeness for the concatenation methodology (blue) versus the complete dendritic arbor (gray). **(B)** Tracing speeds for various neurons in the lateral horn and gnathal ganglion, traced to lower levels of completeness primarily for cell type identification (mean path length for different completeness indicated for reference). Cells are grouped by reconstruction methodology as in (A). Linear fits indicate tracing speed-ups from using FFN concatenation of 5.4x (A) and 14.6x (B).

**Figure 7.**
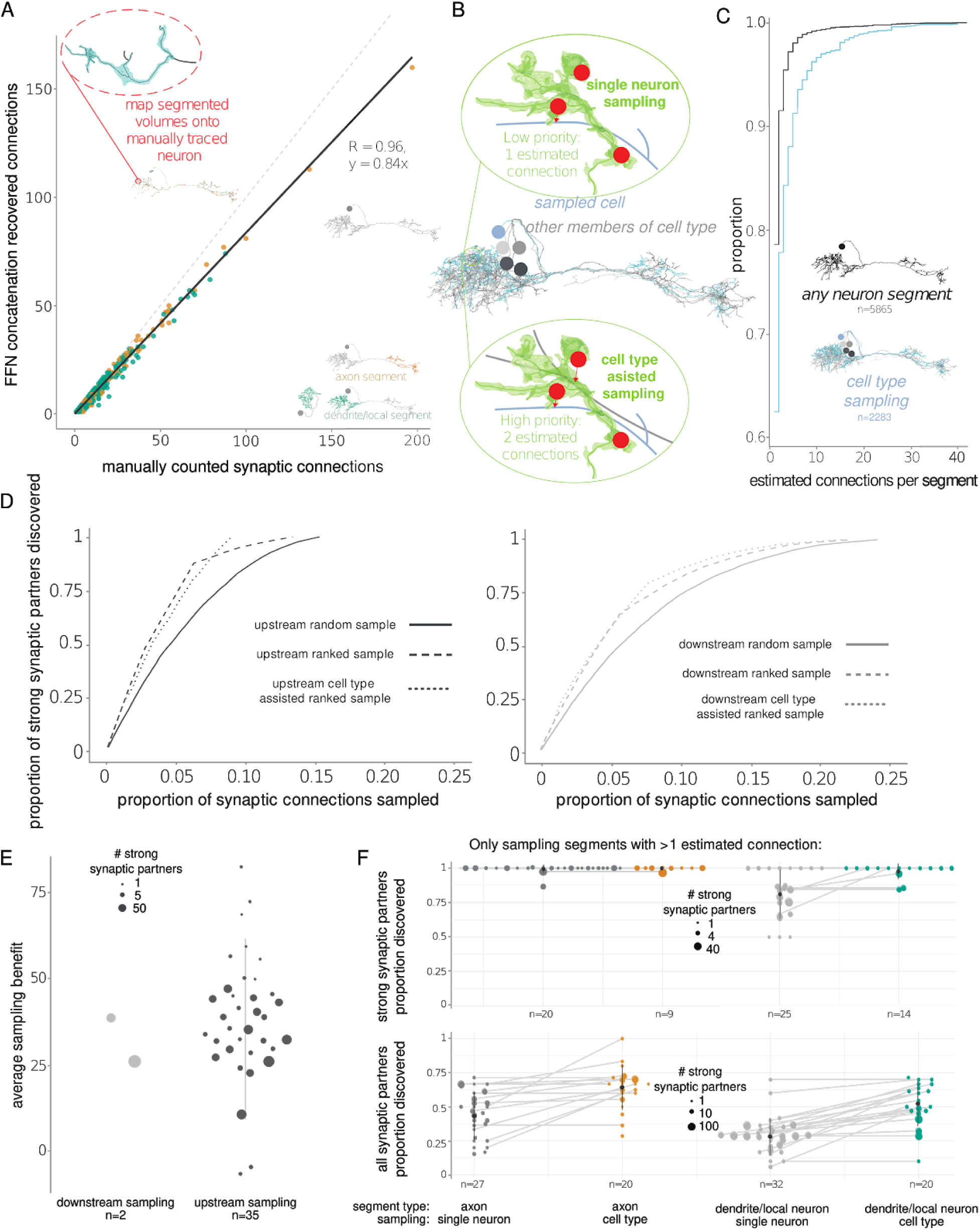
Trans-synaptic circuit tracing. **(A)** For a set of manually traced neurons in the lateral horn (LH, n=82), the number of incoming synapses recovered per upstream partner by concatenating FFN segments, versus ground truth synapse total from manual search. Upstream partners have either axonal morphology (orange points) or dendritic morphology consistent with a local interneuron (green points). Points along the dashed gray unity line indicate near-perfect synapse recovery. The linear fit indicates 84% recovery on average. **(B)** Illustration of two ways to use FFN segments to recover strong upstream synaptic partners. For a traced neuron of interest (blue), we use the FFN segments (top) to search for clusters of marked incoming synapses (red spheres) that originate from a single upstream partner; these are probable strong connections. If other cells of the same type have also been traced (gray), we can also identify FFN segments containing multiple synapses onto any cell of the type of interest as probable strong partners (bottom). **(C)** Cumulative density plot for the proportion of upstream fragments discovered, and the number of connections they have with the given neuron. FFN segmentation-assisted sampling upstream of single starter neurons (black, n=82) and sampling by cell type (blue, n=54 neurons that have >1 neuron in their cell type traced). **(D)** Example average sampling curves for finding strongly connected partners (connected by five or more synapses) of LH neurons. Left, sampling curves for downstream partners of a DA2 projection neuron (Huoviala et al. 2018). We assessed the recovery of strongly connected synaptic partners with different synapse sampling strategies (n=100); solid line: completely random sampling; dashed line: a sample of downstream FFN segments ranked by the number of estimated connections (“ranked sampling”); dotted line: a sample of downstream FFN segments ranked by the number of estimated connections from all five members of this cell type (“cell type assisted ranked sampling”). Right, sampling curves for upstream partners of a PD2a1 LH output neuron (Dolan et al. 2018). Again, five neurons of the same cell type were used for cell type assisted ranked sampling. **(E)** Tracing benefit, i.e. the average proportion of randomly sampled synapses from which one would no longer need to reconstruct (to discover the same number of strong synaptic partners) if using the ranked sampling approach. **(F)** Top, swarm plots of the proportion of strongly connected upstream partners that can be recovered by sampling only from FFN segmented fragments that contain two or more connections to the neuron of interest (gray) or to any neurons of the same cell type (color). Bottom, proportion of all partners, strong or weak. Left, neurons whose upstream segment has axonic morphology. Right, upstream segments with dendritic morphology, consistent with local interneurons.

As part of an ongoing effort to characterize the network connectivity of the mushroom body calyx (Zheng et al. 2020), we traced the dendritic arbors of 1,146 Kenyon cells (KCs) to determine the complement of olfactory projection neuron (OPN) inputs to each (Fig. 6A). All tracing was done in the CATMAID environment (Saalfeld *et al*., Schneider-Mizell *et al*.) (see Methods). An initial group of 545 KCs was traced entirely via manual node marking, with each arbor reconstructed completely. For a second group of 601 KCs, the arbor was instead reconstructed by manually concatenating segmentation-derived skeleton fragments, with limited manual node marking to link between automated skeletons as needed. In this case, the arbor was traced only to the completeness necessary to positively identify all OPN inputs (Fig. 6A, inset). FFN concatenation is a feasible approach in part due to the very low merge error rate in the segmentation (Fig. 5B) as well as the high coverage of the larger (≥ 1 μm path length) automated skeleton fragments that tracers find convenient to work with (Fig. 2B-D). Comparing tracing speeds (person-minutes per μm path length) shows a 5.4x speed-up on average from using FFN skeletons.

In a separate tracing effort, a variety of neurons in the lateral horn (Frechter et al. 2018) and gnathal ganglion (Hsu and Bhandawat 2016) were also reconstructed either manually or via FFN concatenation (Fig. 6B). In this case, the goal of tracing was primarily to allow cell type identification; this generally requires lower levels of completeness, e.g. just the cell body fiber and the major axonal and dendritic branches. Comparing tracing speeds for this workflow shows an 14.6x speed-up on average from using FFN skeletons.

Another common biological analysis workflow is trans-synaptic tracing. In this setting, the goal is to determine for a given neuron how its input and output synapses are distributed, and what the complement of upstream and downstream synaptic partner neurons are. Often there is a particular emphasis on identifying strong partners, e.g. those making five or more synaptic connections with the given neuron.

We assessed the benefits of using FFN segmentation to assist trans-synaptic tracing (Fig. 7). For these experiments, we assumed that a neuron of interest had already been traced, whether manually or via FFN concatenation (Fig. 6), but its upstream and downstream partner neurons had not. We further assumed that all input and output synapse locations had been marked; in this case marking was done manually, but an automated approach (Kreshuk et al. 2015; Buhmann et al. 2018; Heinrich et al. 2018; Huang, Scheffer, and Plaza 2018; Buhmann et al. 2019) would also be viable.

First, we asked how many input synapses onto a given neuron are associated with non-trivial (overlapping at least 3 consecutive manually traced skeleton nodes) upstream FFN segments. Starting from 82 manually traced neurons in the lateral horn, we found that on average 84% of upstream synaptic connections could be recovered by concatenating FFN segments (Fig. 7A). For a number of traced partners recovery was close to 100%, and in some cases synapses were found in the FFN segmentation that had been missed in manual tracing.

Availability of FFN segmentation also allowed more efficient, targeted sampling for strong synaptic partners (Fig. 7B-F). The baseline workflow for strong partner discovery is simply to sample input/output synapses for a given neuron in arbitrary order, and reconstruct each corresponding upstream/downstream partner iteratively to find those making multiple synapses onto the neuron of interest, which implies greater significance to the neural circuit (Frechter et al. 2018). However, if an existing automated segment connects multiple synapses together, then that partner can immediately be prioritized for reconstruction as a likely strong partner. This can be extended if the neuron of interest is one of a class of neurons that have also already been traced; in this case an upstream segment that contains multiple synapses onto the neuron of interest *or any other neurons of the same class* is a likely strong partner (Fig. 7B-D). Segmentation-assisted ranked sampling allowed faster recovery of strong connections for a given level of reconstruction effort, for both upstream and downstream partners and especially for partner axons (Fig. 7D-F). In sum, this allowed human tracers to skip as many as 80% of FFN fragments that were not multiply connected (Fig. 7C) to the neuron of interest and still recover the large majority of strongly connected partners (median 97.6% recovered for cell type ranked sampling with axonic partners; 100% for dendritic partners; Fig. 7F, top).

## Discussion

We have demonstrated that it is possible to segment a complete ssTEM volume of a *Drosophila* brain using a combination of Flood-Filling Networks and novel procedures that compensate for imperfections in the raw data and its initial global alignment. To our knowledge, FAFB-FFN1 is the largest dense segmentation of a central nervous system that has been publicly released, and the first “whole brain” dataset that has been densely segmented by computational means.

The approaches introduced here handle data irregularities explicitly, detecting them and preprocessing on the fly to correct them before the segmentation model sees the data. An alternative approach is to train the segmentation model to take irregularities in stride, e.g. via training data augmentations that simulate the spectrum of issues present in the dataset (K. Lee et al. 2017). We found that while including these augmentations during FFN training improved segmentation robustness, it was not sufficient to allow competitive segmentation of FAFB without explicit handling of irregularities.

FAFB-FFN1 is largely an “over-segmentation” in which each neuron in the automated segmentation is composed of a number of separate segments, each of which average 199 μm in path (run) length when sampled uniformly. Experiments with human tracers have shown that it is more efficient, by an order of magnitude, for humans to manually concatenate automatically-generated segments, as compared to performing manual skeletonization. Moreover, some highly useful biological inferences, such as identification of neuronal cell type, can be performed using FAFB-FFN1 in minutes or seconds per neuron, as compared to hours in the purely manual case. Experiments also demonstrated the applicability of FAFB-FFN1 to circuit tracing and connectomic analyses; these studies combine segmentation with an accounting of the synaptic connections between neurons, for which large-scale automated methods on *Drosophila* tissue are an active research area (Kreshuk et al. 2015; Buhmann et al. 2018; Heinrich et al. 2018; Huang, Scheffer, and Plaza 2018; Buhmann et al. 2019).

Further improvements in the automated reconstruction of volumes such as FAFB may be driven by advances in segmentation methodology. However, a survey of errors in FAFB-FFN1 that persist even after preprocessing for data irregularities suggests that further improvement to global or local alignment methods could improve results even with the same segmentation approach. Furthermore, the FFNs used to generate FAFB-FFN1 were trained primarily on a small subset of the volume located in one particular region of the fly brain (Funke et al. 2016), augmented with a handful of sparsely proof-read neurons sampled from other regions. Training on additional ground truth that represents greater coverage of distinct regions and cell types in the fly brain may also improve reconstruction accuracy. One alternative segmentation of FAFB based on further refinement of the global alignment and an expanded training dataset has recently been released through the open-access, centrally-managed FlyWire tracing environment (Dorkenwald et al. 2020). Although not free to download, this resource will further enhance circuit analyses. Given that error modes between FAFB-FFN1 and FlyWire are frequently uncorrelated, there is also an opportunity to combine both resources in tracing workflows, provided their differences in global alignment and data accessibility can be reconciled.

In addition to accelerating the reconstruction and study of *Drosophila* neurons and circuits, FAFB-FFN1 may provide a useful resource for the development of novel computational tools. For example, there is a largely unexplored opportunity to develop novel segment agglomeration algorithms that exploit truly brain-wide neuron shape representations and priors. Large-scale morphometric analysis of *Drosophila* neurites and neuron shapes (Sümbül et al. 2014; Zhao and Plaza 2014; Costa et al. 2016; Kanari et al. 2018; Chandrasekhar and Navlakha 2019), along with efforts to model detailed biophysics, such as second messenger dynamics along neurites (Rangamani et al. 2019), may also immediately benefit from sampling segments within FAFB-FFN1.

## Methods

### Input and training data

The raw image and training data were all derived from the Full Adult Fly Brain (FAFB) dataset described by (Zheng et al. 2018). The dataset was acquired at 4×4×40 nm nominal resolution, while our segmentation pipelines additionally used volumes downsampled to 8×8×40, 16×16×40, 32×32×40, and 64×64×40 nm via non-overlapping boxcar-mean filtering. Prior to training and inference it was helpful to normalize all raw imagery via Contrast Limited Adaptive Histogram Equalization (CLAHE) (Zuiderveld 1994), with kernel sizes of 2048×2048 nm followed by 1120×1120 nm.

### Evaluation data and metrics

To evaluate automated segmentation quality, we compared our results to manually-traced ground-truth neuronal skeletons (Fig. 2). We used the set of 166 ground-truth skeletons from the mushroom body region, described in (Zheng et al. 2018), as well as a partially overlapping larger collection of 405 Kenyon cells and 123 olfactory projection neurons (Zheng et al. 2020). Where we list IDs for specific FAFB neurons (Fig. 2, and below), they are taken from the FAFB public skeleton IDs (Zheng et al. 2018). We further quantified the agreement between automated segments and ground-truth skeletons via *skeleton metrics*. As described previously (Michał Januszewski et al. 2018), these metrics consist of *edge accuracies* and *expected run lengths* (ERLs).

Briefly, *edge accuracies* count the proportion of ground-truth skeleton edges falling into four non-overlapping categories: correct edges, whose ends are both within one segment; merge errors, where either end is within a segment that erroneously joins two ground truth skeletons; split errors, whose ends are in two different merge-free segments; and omission errors, where either end is in an unsegmented area. ERL computes the expected error free path length (the linear distance connected by correct skeleton edges) in the segmentation given a uniformly sampled starting position along a ground-truth skeleton.

The training data consisted of 3 densely labeled cutouts from the Mushroom Body region of an earlier FAFB global alignment, provided by the MICCAI Challenge on Circuit Reconstruction from Electron Microscopy Images (CREMI) (Funke et al. 2016). Each cutout totals 1250×1250×125 labeled voxels at 4×4×40 nm. We downsampled these cutouts to train networks at reduced resolutions. To generate additional training data at 16×16×40 nm resolution, we additionally proofread 201.3 megavoxels of an earlier segmentation result to rough topological completion. For segmentation pipeline development (Figs. 3, 4) we also used a larger (120,000 × 120,000 × 75,040 nm) unlabeled cutout, referred to as “Sample E”, from around the mushroom body of the v14 alignment (starting offset 376,000 × 80,000 × 158,200 nm).

It was helpful to erode the ground truth skeletons by 1 node back from all branch endpoints, due to many cases where the manually placed endpoint nodes were directly on the border between neighboring cells. We also ignored merge errors that involved fewer than 3 ground truth skeleton nodes (Berning, Boergens, and Helmstaedter 2015). These smallest detected mergers often reflected errors in ground truth skeleton node placement, or were expected due to irregular section substitution changing the positions of neurons within substituted sections, or else were real segmentation errors but were too minor to impact biological interpretation or subsequent processing. After inspecting a subset of manual tracings, we also removed 169 nodes that were reported as larger merge errors but were found to be misplaced, as well as all nodes occurring in sections 4411 and 4423, which were subject to irregular section substitution across most of the XY extent of the manual tracings.

In a handful of exceptional cases we also excluded entire ground truth neurons from evaluations. Olfactory projection neurons 30791 and 51886 were found to be legitimately fused together over about 25 um within the medial antenna lobe tract (mALT). This apparent biological aberration or tissue preparation artifact was reported as a merge error if included in the metrics. Olfactory projection neuron 28876 had abnormally dark cytoplasm, possibly due to cell damage (Fiala, Spacek, and Harris 2002), leading to many split and omission errors as well as a few mergers.

### Misaligned and irregular section detection

Misalignments and irregular data regions were detected via section-to-section cross-correlation template matching (Lewis 1995). Overlapping image patches of about 4×4 μm were extracted at 32×32 nm resolution and 128×128 nm stride across each section, and cross-correlated against patches from the following section. For each image pair, the offset of the peak of the correlation surface was taken as the local section-to-section shift.

We found that cross-correlations based on simple zero-padded search windows, without extended image context or cumulative sum normalization (Lewis 1995), were more robust to areas with data irregularities or coherent lateral movement of neural processes (e.g. fiber tracts running transverse to the cutting plane). Therefore we simply zeroed the image mean prior to cross-correlation, found the peak of the unnormalized correlation surface, and then post-normalized the peak value by the autocorrelation magnitude to generate a measure of template match quality.

The cross-correlation procedure produces a subsampled, quantized section-to-section flow field for the entire volume. In areas of misalignment or coherent lateral movement of neural processes, the flow field reveals the magnitude of local section-to-section shifts. It also detects irregular data areas such as tissue cracks and folds by the characteristic sharp discontinuities in their flow fields, and damaged or occluded sections by their large, inconsistent shift values and low template match quality values.

### Misaligned and irregular section handling

Movement restriction (Michał Januszewski et al. 2018) blocks the FFN during inference from moving through or evaluating on any field of view where flow field shift magnitudes exceed threshold. For FAFB segmentation we set relatively tolerant thresholds of 64-128 nm, but this still resulted in significant movement restriction over the volume. Restriction generally causes local split errors, but when two restricted areas are close to each other it can prevent an area from being segmented at all (Fig. 4A) and thus also causes omitted edge errors.

Local realignment (LR) instead attempts to allow FFN evaluation through misaligned areas by dynamically correcting alignment at inference time (Fig. 3). Before the FFN considers each local subvolume (generally 400×400×100 voxels, or 60×60×30 voxels for agglomeration) we compute a translation-only realignment transform based on the weighted median flow field shift over each section, with cross-correlation match quality used as weights. Excessive section-to-section shifts usually reflect data irregularities rather than misalignments, so we set a threshold of 128-256 nm, above which we discard the shift for a given section, resetting it to zero. Applying the computed realignment transform to the subvolume results in a view of the data in which errors in the global alignment of the input are nominally corrected. The FFN operates on the corrected data, and we then reverse the transform to return the segmentation output back to the global coordinate space.

An important requirement is that the subvolume provided to the FFN after realignment should be cropped to a rectilinear shape containing only valid data, so that the FFN is afforded free movement and has valid context. Similarly, the reverse transformed segmentation output should be cropped rectilinear and valid, so that output subvolumes can be reassembled into a coherent segmentation volume. To compensate for cropping, each subvolume has its bounds expanded before the forward realignment is applied, with the amount of additional context needing to be drawn from the input volume dependent on the largest accumulated XY shifts in the given transform.

Following realignment, the flow field of the subvolume is also recomputed, so that any large remaining shifts can still trigger FFN movement restriction. This is important in cases where highly distorted, cracked, or folded sections can only be adequately realigned over a portion of the subvolume XY extent, as well as for data irregularities that realignment cannot address such as missing or occluded sections. In preliminary experiments, we found that replacing translation-only LR with an affine alignment approach could successfully address some of these areas. However, affine LR proved more difficult to regularize.

For data issues inadequately addressed by LR, irregular section substitution (ISS) attempts to allow FFN evaluation through the area by replacing the affected section locally within the subvolume with data from a neighboring section (Fig. 4). The criterion for considering a section for substitution is whether the magnitude of flow field shifts for the unsubstituted section triggers significant movement restriction (defined as restriction over more than 3% of the section extent within the subvolume). If so, the section is replaced with the preceding section, and the cross-correlation flow field with respect to the following section is recomputed. If the new flow field results in a significant reduction in movement restriction (defined as 50% or more reduction), the candidate substitution is accepted and the LR transform is updated to reflect the new flow field.

Thus ISS is only allowed when the sections on either side of an irregularity can be adequately realigned to each other. The anisotropy of the FAFB dataset makes it rare for this to be effective across more than a single consecutive section. For FAFB-FFN1, we considered only single section substitutions, except for the 32×32×40 nm agglomeration stage, where we allowed up to 3 consecutive sections to be substituted.

### Flood-filling network training and inference

FFNs were trained as described previously (Michał Januszewski et al. 2018) and as publicly released (Michal Januszewski 2019). Briefly, a deep 3d convolutional neural network was trained in TensorFlow via asynchronous stochastic gradient descent. The network field of view was 33×33×17 voxels except networks trained at full 4×4×40 nm resolution, where the field of view was 33×33×9. The network architecture consisted of a series of 18 convolutional layers with 3×3×3 kernels and 32 feature channels, paired off into 9 residual units with no downsampling, resulting in 472,353 trainable parameters. All three CREMI cutouts of ground-truth labels were used for training, downsampled to the appropriate resolution as needed.

FFN seeding and movement policies for training and inference were as described previously. Due to the anisotropy of serial-section data, we also found that a 2d variant of the Sobel-Feldman peak distance seeding procedure (Michał Januszewski et al. 2018) was effective for filling in small processes in later inference runs. FFN inference hyperparameters were set similarly to previously described: initial field-of-view fill value 0.05; movement threshold 0.9; segment threshold 0.6. The base movement step size was 8×8×4 voxels.

Network weights were checkpointed periodically during training, and convergence was assessed at each checkpoint by running test inference on a small cutout from the mushroom body calyx (“calyx1”) and evaluating skeleton metrics against the resulting segmentation. We then selected a subset of the checkpoints whose calyx1 evaluations had zero merge errors and high overall edge accuracy for inference over the larger Sample E mushroom body cutout. Finally, we selected the checkpoints with the lowest merge error rates and highest overall edge accuracies from these evaluations for inference over all of FAFB (Fig. 5). Two additional checkpoints were selected for oversegmentation consensus (Michał Januszewski et al. 2018) at 16×16×40 nm by reevaluating all checkpoints over an additional small area (“adaptive1”) where the base segmentation had remaining merge errors, and selecting checkpoints that corrected these errors while maintaining high edge accuracy.

### Segmentation-Enhanced CycleGAN for Triple-Section Interpolation

A SECGAN (Michal Januszewski and Jain 2019) was trained to synthesize missing data in the raw FAFB volume at 16×16×40 nm voxel resolution. Both “input” dataset X and “target” dataset Y were sampled from regions within FAFB. Samples for X were chosen from regions containing data with no known irregularities, and samples from Y were specifically chosen within regions that contained three consecutive missing or irregular sections. The field of view of the generator was 33×33×33 voxels, using a ResNet-like architecture previously described (Michal Januszewski and Jain 2019), and a ResNet18 discriminator architecture (He et al. 2016). During training, the central 3 sections of every training example were excluded from the Y cycle loss, and were zeroed-out in the input of the Y discriminator. The Y images were also altered by filling the empty sections with the contents of the directly preceding non-empty section.

For SECGAN inference, the raw image data around the missing sections was first elastically realigned (Saalfeld et al. 2012) via an iterative procedure. First, the flow field between the two sections directly preceding and following the gap was estimated from downsampled imagery at 64×64 nm in-plane pixel resolution using patch-wise cross-correlation (patch size 160 pixels, stride 40 pixels), and used to relax a 2-section (ignoring the gap) elastic mesh. The procedure was then repeated at 32 × 32 nm in-plane resolution to obtain the final alignment.

After running FFN segmentation within 200-section subvolumes centered on the SECGAN substituted region, overlaps were computed between the SECGAN segments and the preexisting segments on either side of the substituted gap. For each SECGAN segment, the maximally overlapping pre- and post-gap preexisting segments were determined, and these two segment IDs were then joined if both overlaps exceeded 1000 voxels. By construction, this approach can only link a single pre-gap segment to a single post-gap segment for each SECGAN segment, which limits the ability to fully address neurites that branch within the substituted block. However, we found this constraint to be important for avoiding new merge errors.

### Overall segmentation pipeline details

The overall pipeline comprised a series of FFN segmentation steps, selected to maximize overall skeleton edge accuracy and ERL while minimizing merge errors, as described previously (Michał Januszewski et al. 2018) and above. For filtering agglomeration decisions, we found that it was more effective to require the proportion of deleted voxels to be below threshold for both segment A and B (operation AND), rather than for either segmentation A or B (operation OR) as done previously.

Some experiments with human tracers (Fig. 6) were carried out using earlier segmentations, referred to as FAFB-FFN0 and SAMPLE-E-FFN0. Detailed hyperparameters for the different pipelines are given in Table 2.

**Table 1.**
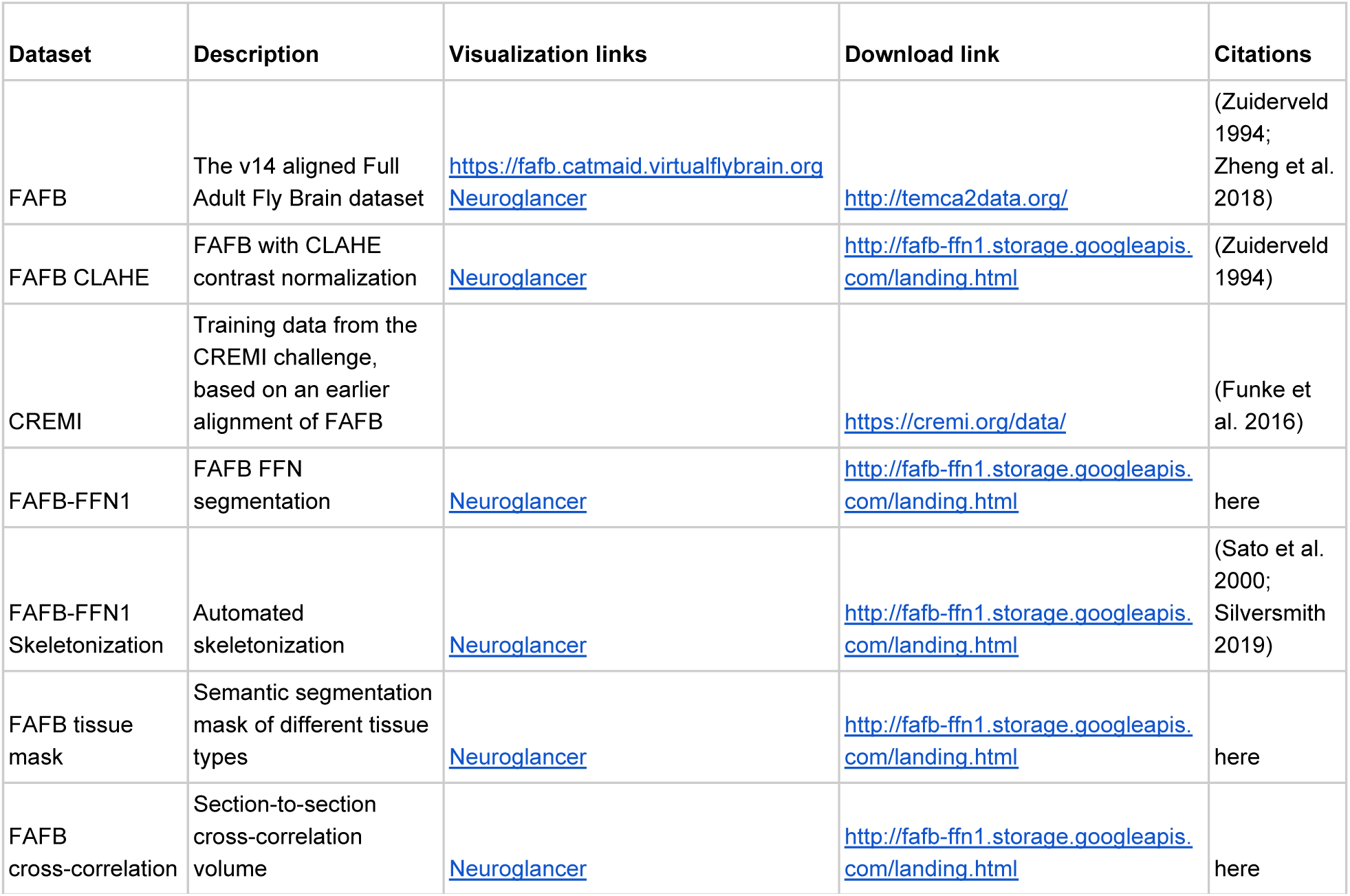
Input data and results.

**Table 2.**
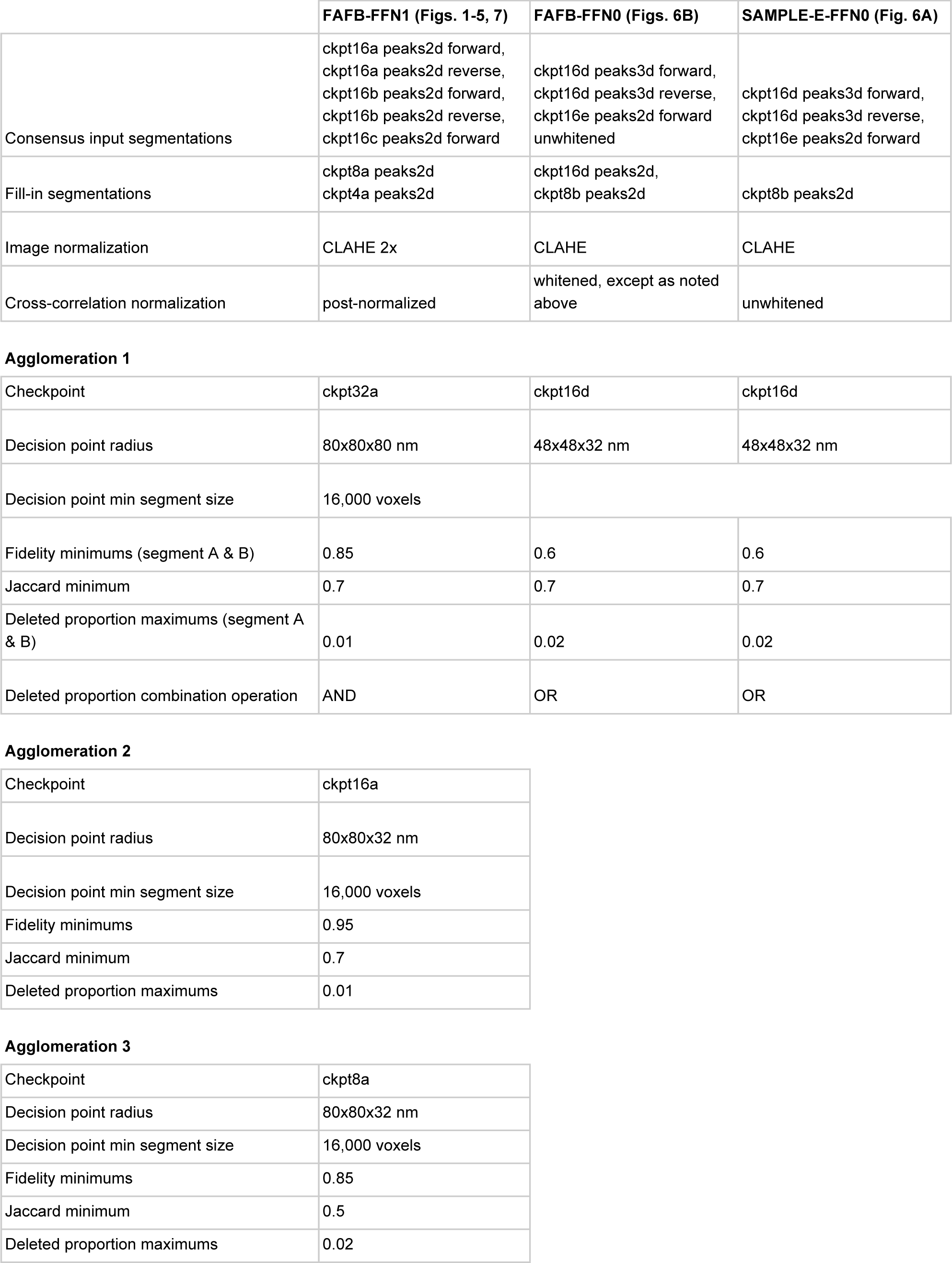
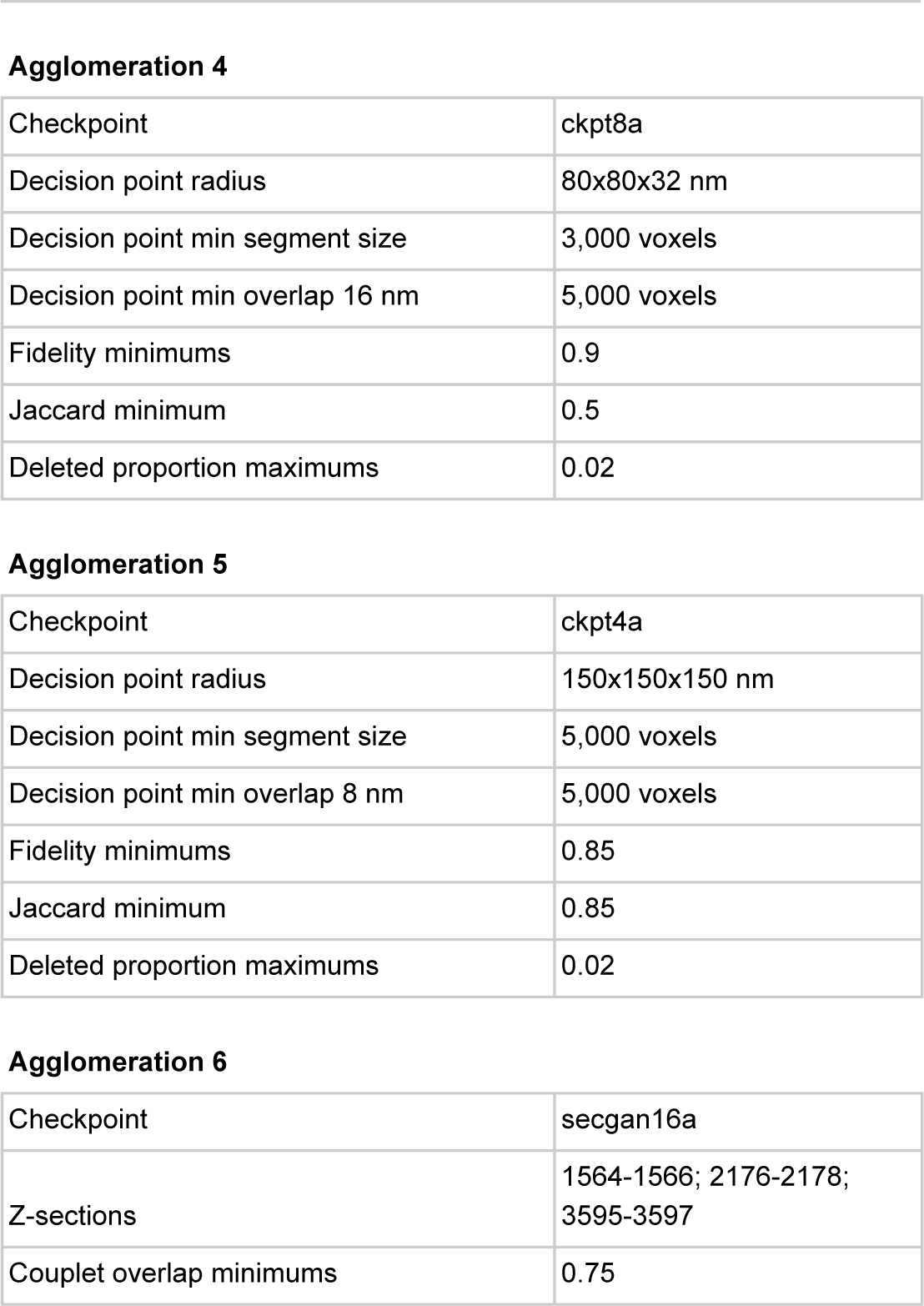
FFN pipeline hyperparameters. All checkpoint names include the XY resolution used in training and inference. peaks3d, 3d Sobel-Feldman peak seeding; peaks2d, 2d Sobel-Feldman peak seeding; forward, normal seed order; reverse, reversed seed order; post-normalized, post-normalized cross-correlation (see Methods); whitened, square-root frequency amplitude normalized image cross-correlation; unwhitened, cumulative sum normalized image cross-correlation

|  | FAFB-FFN1 (Figs. 1-5, 7) | FAFB-FFN0 (Figs. 6B) | SAMPLE-E-FFN0 (Fig. 6A) |
| --- | --- | --- | --- |
| Consensus input segmentations | ckpt16a peaks2d forward,<br>ckpt16a peaks2d reverse,<br>ckpt16b peaks2d forward,<br>ckpt16b peaks2d reverse,<br>ckpt16c peaks2d forward | ckpt16d peaks3d forward,<br>ckpt16d peaks3d reverse,<br>ckpt16e peaks2d forward<br>unwhitened | ckpt16d peaks3d forward,<br>ckpt16d peaks3d reverse,<br>ckpt16e peaks2d forward |
| Fill-in segmentations | ckpt8a peaks2d<br>ckpt4a peaks2d | ckpt16d peaks2d,<br>ckpt8b peaks2d | ckpt8b peaks2d |
| Image normalization | CLAHE 2x | CLAHE | CLAHE |
| Cross-correlation normalization | post-normalized | whitened, except as noted<br>above | unwhitened |
| Agglomeration 1 |  |  |  |
| Checkpoint | ckpt32a | ckpt16d | ckpt16d |
| Decision point radius | 80x80x80 nm | 48x48x32 nm | 48x48x32 nm |
| Decision point min segment size | 16,000 voxels |  |  |
| Fidelity minimums (segment A & B) | 0.85 | 0.6 | 0.6 |
| Jaccard minimum | 0.7 | 0.7 | 0.7 |
| Deleted proportion maximums (segment A & B) | 0.01 | 0.02 | 0.02 |
| Deleted proportion combination operation | AND | OR | OR |

Agglomeration 2
|  |  |
| --- | --- |
| Checkpoint | ckpt16a |
| Decision point radius | 80x80x32 nm |
| Decision point min segment size | 16,000 voxels |
| Fidelity minimums | 0.95 |
| Jaccard minimum | 0.7 |
| Deleted proportion maximums | 0.01 |

Agglomeration 3
|  |  |
| --- | --- |
| Checkpoint | ckpt8a |
| Decision point radius | 80x80x32 nm |
| Decision point min segment size | 16,000 voxels |
| Fidelity minimums | 0.85 |
| Jaccard minimum | 0.5 |
| Deleted proportion maximums | 0.02 |

Agglomeration 4
|  |  |
| --- | --- |
| Checkpoint | ckpt8a |
| Decision point radius | 80x80x32 nm |
| Decision point min segment size | 3,000 voxels |
| Decision point min overlap 16 nm | 5,000 voxels |
| Fidelity minimums | 0.9 |
| Jaccard minimum | 0.5 |
| Deleted proportion maximums | 0.02 |

Agglomeration 5
|  |  |
| --- | --- |
| Checkpoint | ckpt4a |
| Decision point radius | 150x150x150 nm |
| Decision point min segment size | 5,000 voxels |
| Decision point min overlap 8 nm | 5,000 voxels |
| Fidelity minimums | 0.85 |
| Jaccard minimum | 0.85 |
| Deleted proportion maximums | 0.02 |

**Table 2. FFN pipeline hyperparameters.** All checkpoint names include the XY resolution used in training and inference.
|  |  |
| --- | --- |
| Checkpoint | secgan16a |
| Z-sections | 1564-1566; 2176-2178;<br>3595-3597 |
| Couplet overlap minimums | 0.75 |

### Tissue Masking

We trained a convolutional network to predict whether a voxel belonged to one of six categories that represented general structural features of the image volume. We manually labeled 10.7 million voxels at 2x reduced lateral resolution as either neuropil (4.6M voxels), cell body (1.6M voxels), glia (0.11M voxels), black border (0.7M voxels), resin (1.6M voxels), or tissue border (1.9M voxels). Annotations were sparsely created using a custom web-based tool (“Armitage”) that enabled manual painting of voxels with a modifiable brush size.

We then used TensorFlow to train a 3d convolutional network to classify a 65×65×5 patch centered on each manually labeled voxel. The network contained three “convolution-pooling” modules consisting of convolution (3×3×1 kernel size, 64 feature channels, VALID mode where convolution results are only computed where the image and filter overlap completely) and max pooling (2×2×1 kernel size, 2×2×1 stride, VALID mode), followed by one additional convolution (3×3×1 kernel size, 16 feature maps, VALID), a fully connected layer that combines information from all 5 slices (512 nodes), and a six-class softmax output layer. We trained the network by stochastic gradient descent with a minibatch size of 32 and 6 replicas. During training, each of the six classes was sampled equally often. Training was terminated after 0.5 million updates.

Inference with the trained network was applied to all voxels in the image volume using dilated convolutions, which is several orders of magnitude more efficient than a naive sliding-window inference strategy. Inference on the whole volume at 16×16×40nm resulted in 1.97 teravoxels of predicted neuropil, 0.25 teravoxels of soma, 0.24 teravoxels of glia, 0.24 teravoxels of black border, 0.46 teravoxels of resin, and 0.41 teravoxels of tissue border.

Tissue masking predictions were used during FFN inference to block FFN evaluations centered at locations with less than 12% predicted neuropil probability and less than 50% predicted soma probability. This improved efficiency of segmentation and prevented some segmentation errors in areas such as glia with textures underrepresented in the training set. Class-wise voxel counts from the tissue masking volume were used to estimate the total dataset size of 39.4 teravoxels of combined neuropil, soma, and glial tissue at full 4×4×40 nm resolution. These three classes were also combined to generate a whole brain mesh for visualization (Fig. 1).

### Automated skeletonization

Some biological analysis workflows require a neuron skeleton representation (Fig. 2A-C, Fig. 6A inset) rather than volumetric segments. We used the TEASAR algorithm (Sato et al. 2000) to automatically convert the FFN segmentation to a skeleton representation. Earlier versions of the segmentation were skeletonized via a different pipeline than the final release; the differences are detailed in Table 3.

**Table 3.** Skeletonization parameters

|  | FAFB-FFN1 (Fig. 7) | FAFB-FFN0; SAMPLE-E-FFN0 (Fig. 6) |
| --- | --- | --- |
| Block size | 256x256x256 | 512x512x256 |
| Resolution (nm) | 32x32x40 | 32x32x40 |
| Maximum reconnection distance (nm) | 50 | 165 |
| Max fragments per skeleton | 1,000,000 | 10,000 |
| Endpoint erosion (nm) | 100 | 250 |
| Node sparsification distance (nm) | 300 | 250 |
| Length / diameter minimum ratio | NA | 10 |
| Invalidation radius scale factor | 4 | NA |
| TEASAR implementation | (Silversmith 2019) | (Funke 2017) |

The scale of FAFB required that TEASAR be run block-wise and at downsampled resolution. To reconnect skeletons at block boundaries, a simple heuristic was used to join the nearest neighbor nodes on either side provided their separation distance was below a fixed threshold. Occasionally this heuristic failed to reconnect across blocks, in which case a single skeleton ID in the result might comprise multiple components. In a few cases, large irregular bodies such as glia, trachea, somas, or non-biological material created highly fragmented skeletons; skeletons with total number of components above a fixed threshold were discarded.

Skeletons were further post-processed to enhance suitability for FFN concatenation workflows (Fig. 6). First, we eroded skeletons back from all endpoints. Skeleton branches with a low path length relative to average diameter often reflect thickening or bumps at the surface of neurites, rather than true neuronal branches; these were either pruned off (for length / diameter < 10; earlier releases) or were excluded from skeletonization by increasing the invalidation radius scale factor (FAFB-FFN1 final release). Finally skeletons were sparsified to a target distance between nodes within unbranched sections. After these post-processing steps, all resulting skeletons greater than 1 μm path length were exported to the tracing environment.

### Neuron skeleton tracing and trans-synaptic analysis

All neuron skeleton tracing (Fig. 6) was done in the FAFB CATMAID environment (Saalfeld et al. 2009). CATMAID provides an interface for exploring the EM image volume, and neurons can be traced manually by marking a series of node points (Schneider-Mizell et al. 2016). The TEASAR-generated skeletons of FFN segments were also imported into CATMAID, and tools were provided for linking skeleton fragments together (“FFN concatenation”) and quickly jumping to fragment endpoints to check for missing continuations of neurites.

Neuronal reconstruction in mushroom body calyx (Fig. 6A) was typically done by two team members, an initial tracer and a subsequent proofreader who validated the tracing, potentially sending issues back to the tracer to iterate on. Neurites in the fly brain can be classified into larger, microtubule-containing neurites (“backbones”) and fine, microtubule-free neurites (“twigs”) (Schneider-Mizell et al. 2016). Microtubules were used as guides during manual tracing and FFN concatenation to ensure quick reconstruction of backbones of the neurons, which is often sufficient for cell type identification.

The CATMAID environment automatically records the amount of time spent tracing each neuron. When the FFN concatenation tracing methodology was introduced, there was a ramping-up period while human tracers adjusted and software tools for efficient concatenation matured. Thus, out of 916 total Kenyon cells (KCs) traced via FFN concatenation, the first 315 had an average tracing speed of 20 seconds per μm path length with a standard deviation of 13 seconds, while the remaining 601 KCs from the later mature period averaged 9 seconds per μm with a standard deviation of 5 seconds. In the Results section, we therefore exclude the ramping-up period from the analysis.

Trans-synaptic analysis combined use of both skeletons and volumetric segments. Software tools for mapping between CATMAID skeletons and the FFN segments are available publicly (Jefferis 2018b). To simulate the recovery of strong synaptic partners using different sampling strategies over synaptic connections (Fig. 7D-E), we started from a fully traced neuron of interest. We also traced at least the local arbor and cell body fiber of all upstream or downstream synaptic partners for purposes of simulation. We then picked a single synaptic connection from which to run simulated tracing, sampled either randomly or using FFN segments to rank multiply connected partners higher *a priori*. For each sampled connection, we recorded whether it corresponded to a strong partner (at least 5 synaptic connections). We also marked all other connections for that partner as visited, under the assumption that they would be discovered in tracing the partner to identification, or else would be trivially connected to the partially traced partner if sampled subsequently. Sampling was then repeated until all strong partners were discovered for the neuron of interest, and the entire procedure was repeated for each starter neuron.

## Acknowledgements

This work was supported by core support from the MRC (MC-U105188491) and an ERC Consolidator (649111) grant (to GSXEJ); the Howard Hughes Medical Institute and a National Institutes of Health BRAIN Initiative (1RF1MH120679-01) grant (DB); and a Wellcome Trust Collaborative Award (203261/Z/16/Z to GSXEJ, DB). We thank FAFB tracers J. Ali, S. Calle-Schuler, J. Hsu, N. Masoodpanah, C. Fisher, N. Sharifi, L. Kmecova, J. Ratliff, S. Imtiaz, B. Gorko, A. Boyes, A. John, E. Moore, B. Koppenhaver, P. Ranft, B. Harrison, S. Murthy, A. Haddad, A. Adesina, A. Scott, C. Marlin, E. Wissell, Z. Gillis, S. Ali, G. Allred, S. Waters, A. Scott, L. Marin, S. Mohr, M. Lingelbach, E. Spillman, A. Smith, T. Ngo, B. Gampah, M. Ryan, J. Dunlap, N. Reddy, A. Fischel, M. Pleijzier, A. Sheridan, K. A. Jahjah, A. Edmondson-Stait, R. Parekh, I. Salaris, A. Warner, S. Lauritzen, R. Roberts, L. Bueld, C. Ardekani, J. Gonzales, R. Rais, W. Chen, N. Ribeiro, and T. Neves for their work reconstructing neurons in mushroom body. We thank I. Varela, B. Morris, S. N. Bailey, and R. J. V. Roberts for their tracing work in lateral horn and gnathal ganglion. We thank T. L. Dean, A. Pope, N. Apthorpe, and J. Shlens for technical contributions and discussions.

